# Diffusion MRI reveals in vivo and non-invasively changes in astrocyte function induced by an aquaporin-4 inhibitor

**DOI:** 10.1101/2020.02.13.947291

**Authors:** Clément S. Debaker, Boucif Djemai, Luisa Ciobanu, Tomokazu Tsurugizawa, Denis Le Bihan

## Abstract

The Glymphatic System (GS) has been proposed as a mechanism to clear brain tissue from waste. Its dysfunction might lead to several brain pathologies, including the Alzheimer’s disease. A key component of the GS and brain tissue water circulation is the astrocyte which is regulated by acquaporin4 (AQP4), a membrane-bound water channel on the astrocytic end-feet. Here we investigated the potential of diffusion MRI to monitor astrocyte activity in a mouse brain model through the inhibition of AQP4 channels with TGN-020. Upon TGN-020 injection, we observed a significant decrease in the Sindex, a diffusion marker of tissue microstructure, and a significant increase of the water diffusion coefficient (sADC) in cerebral cortex and hippocampus compared to saline injection. These results indicate the suitability of diffusion MRI to monitor astrocytic activity in vivo and non-invasively.

## Introduction

Proper neuronal function necessitates a highly regulated extracellular environment in the brain. Accumulation of interstitial solutes, such as amyloid β and toxic compounds [1] may lead to degenerative diseases, such as Alzheimer’s disease [2,3] or even autism [4]. Although the Blood Brain Barrier is thought to be the primary mechanism involved in controlling the brain blood-exchange, the existence of a fluid driven transport system (so-called glymphatic system) via cerebrospinal fluid (CSF) or interstitial fluid (ISF) has been proposed recently as a waste clearance system through the perivascular and interstitial spaces in the brain [1,5]. Hypothetically, CSF crosses the astrocyte end-feet bound to arteries in the perivascular space [6,7]. After washing the interstitial space, the resulting ISF is flushed back outside the brain via veins in the perivascular space. Several factors play a crucial role in the modulation of this clearance system activity, notably sleep and anesthesia, perhaps via a modulation of brain blood volume and pressure [8–11]. This scheme gives astrocytes a crucial role in controlling water movements between the blood and the brain, through a mechanism dependent on Aquaporin-4 (AQP4), a membrane-bound water channel expressed at their end-feet [12].

Deletion of the AQP4 gene suppresses the clearance of soluble amyloid β [1]. As a consequence of the regulation of astrocyte membrane water permeability, AQP4 is involved in the rapid volume regulation of astrocytes [13]. Although most studies were performed *in vitro*, some studies showed astrocyte volume change *in vivo* using 2-photon microscopy. Acute osmotic and ischemic stress induce astrocyte volume changes *in vivo* mice [14], and Thrane et al showed that the astrocyte volume change induced by osmotic stimulation was inhibited in AQP4 KO mice [15]. Overall, such studies suggest that dynamic volume change of astrocytes may be associated with CSF flow regulation. Actually, astrocytes end-feet are involved in the CSF/ISF exchanges in perivascular space during sleep/awake cycle [16].

This clearance system has been evidenced initially using fluorescent tracers and 2-photon microscopy in rodents [1], however MRI offers a more versatile approach. MRI studies have relied on the intrathecal or intravenous injections of gadolinium-based contrast agents (GBCAs) as tracers [7,17].

However, this approach remains invasive and, paradoxically, gadolinium has been shown to deposit in the brain [18,19] possibly in relation to a lack of brain drainage [10]. Therefore, alternative methods are needed to investigate the glymphatic system non-invasively, especially in the human brain. Fluid-dynamics driven and BOLD fast MRI have the potential to evaluate CSF pulsations in the ventricles and hemodynamics [11,20,21], while IVIM and diffusion MRI have been shown as promising methods for the evaluation of the ISF [7,22–24].

Those considerations led us to investigate whether diffusion MRI was sensitive to astrocyte activity and, in turn, could become a marker of the overall glymphatic system. Diffusion MRI is exquisitely sensitive to changes in tissue microstructure, notably cell swelling [25]. Diffusion MRI is, for instance, sensitive to astrocyte swelling induced in rodents [26]. Hence, we hypothesized that dynamic changes in astrocytes activity and related volume changes could be monitored directly and non-invasively with diffusion MRI. To test this hypothesis we monitored variations of new diffusion MRI markers, namely the Sindex and the sADC, which have been tailored to increase sensitivity to tissue microstructure through water diffusion hindrance [27,28] upon acute inhibition of astrocyte AQP4 channels in a mouse brain model. To do so we used 2-(nicotinamide)-1,3,4-thia-diazole (TGN-020), a compound that blocks AQP4 channels *in vivo* in the mouse brain [29].

## Material and methods

### Animal preparation

Thirty-two male C57BL6 mice (16-28 g, Charles River, France) were allocated to two groups. First, for the TGN-020 group, 16 mice received an intra-peritoneal injection of 250mg/kg TGN-020 diluted in a gamma-cyclodextrine solution (10 mM) in order to increase its solubility. Second, for the control group, sixteen mice received an intra-peritoneal injection of the vehicle solution only, 10 mM gamma-cyclodextrine in saline. The mice were housed on a 12-hour light-dark cycle and fed standard food ad libitum. Anesthesia was induced using 3% isoflurane in a mix of air and oxygen (air: 2 L/min, O2: 0.5 L/min). Then, 0.015 mg/kg of dexmedetomidine was administered intraperitoneally and followed by a continuous infusion of 0.015 mg/kg/h via subcutaneous catheter and maintained isoflurane at 0.8%.

Throughout the acquisition, the animals’ body temperature was maintained between 36.5 and 37.0 °C using heated water (Grant TC120, Grant Instruments, Shepreth, UK). To avoid motion-related artifacts the head was immobilized using a bite bar and ear pins. The respiration rate was monitored and stable (60-90 /min) throughout the experiment.

All animal procedures used in the present study were approved by an institutional Ethic Committee and government regulatory agency (reference APAFIS#8462-2017010915542122v2) and were conducted in strict accordance with the recommendations and guidelines of the European Union (Directive 2010/63/EU). This manuscript is in compliance with the ARRIVE guidelines (Animal Research: Reporting in Vivo Experiments) on how to REPORT animal experiments.

### MRI experiments

The MRI experiments were conducted on a Bruker 11.7T scanner (Bruker BioSpin, Ettlingen, Germany) equipped with a gradient system allowing a maximum gradient strength of 760 mT/m. A cryo-cooled mouse brain coil was used. Animal positioning was performed using multi-slice fast low angle shot imaging (FLASH, TE/TR = 2.3/120 ms). A global first and second order shim was achieved followed by a local second order shim over the brain parenchyma. Structural (anatomical) images were acquired with the following parameters: T2 TurboRARE sequence, TE/TR = 11.15/2500 ms. Diffusion-weighted echo-planar imaging (DW-EPI) data sets were acquired with the following parameters: 150×150×250 μm^3^ resolution, 3 b-values (0, 250, 1750) s/mm^2^ along 6 directions, NA = 4, TE/TR = 36.3/2300 ms, 18 slices, diffusion time = 24 ms, total scan time = 5 min. The area covered by the 18 slices of the scans encompassed 5 mm in the axial plane with the center of the slab positioned at the middle of the brain. Six DW-EPI sets were first acquired (baseline) before TGN-020 or vehicle was injected. Then, after the injection, 12 additional DW-EPI data sets were acquired (Fig. 1) every 5 minutes.

**Fig. 1.**
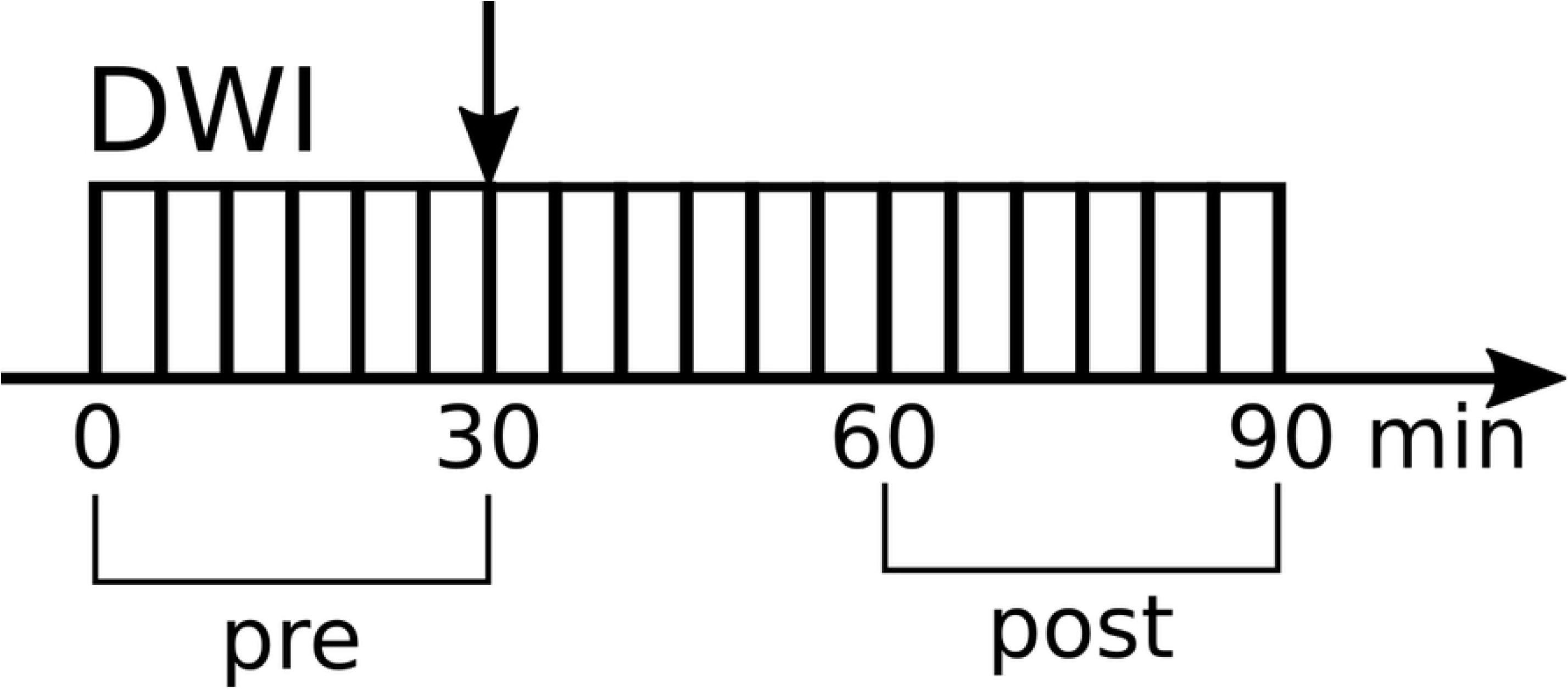
Schematic figure of MRI experiment protocol. The vertical arrow indicates the saline or TGN-020 injection. DWI, diffusion weighted MRI; NR, number of repetition. Horizontal arrow indicates the timeline.

### Data processing

#### Preprocessing

Regions of interest (ROIs) were originally created using the Allen Brain Atlas [30] and registered to a mouse brain MRI template [31] to create a labeled atlas. The atlas was then co-registered with the b=0 images of the first scans for each subject, and not the other-way around to avoid changing raw voxel signals. Those preprocessing steps, also including radiofrequency bias field correction [32] and denoising were performed using ANTs (https://stnava.github.io/ANTs/) [33] (Fig. 2). We assumed no change of position among the different scans during acquisitions for each mouse and geometric distortion with b values to be negligible; this condition was qualitatively checked on several mice. Furthermore, signal instabilities were quantitatively evaluated for each subject and subjects exhibiting instabilities above 4% for most voxels were eliminated. Dataset signals were further subject to noise floor correction [34].

**Fig. 2.**
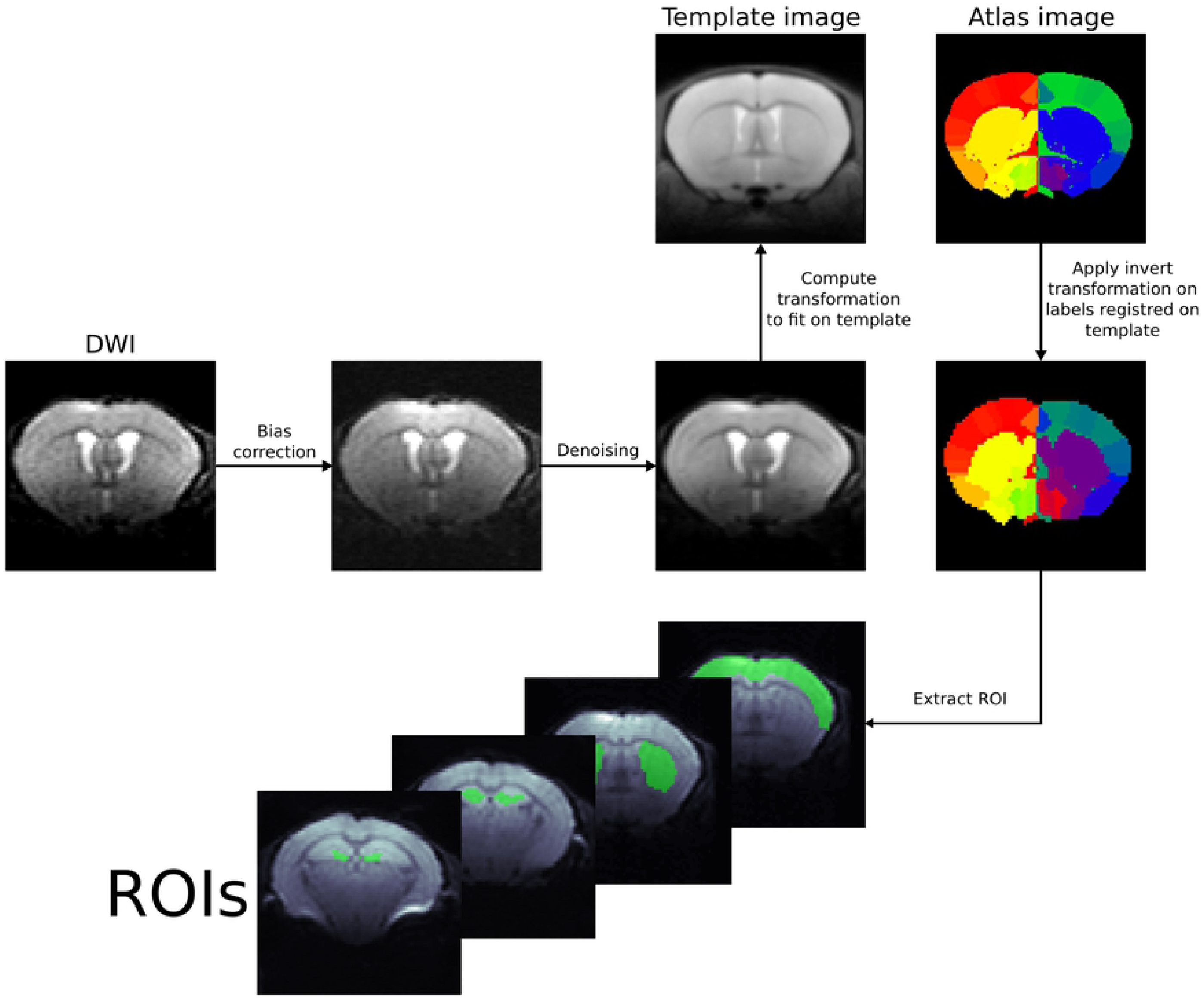
Example of ROI registration with ANTs on a mouse brain.

#### DWI analysis

First, a shifted ADC (sADC) was computed using signal acquired at two key b values (instead of the standard values of 0 and 1000s/mm^2^), chosen to optimize signal sensitivity to both Gaussian and non-Gaussian diffusion which is more sensitive to tissue microstructure through hindrance effects [27,28].

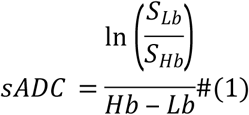

where Lb is the low key b value (250s/mm^2^), Hb the high key b value (1750 s/mm^2^), S_Lb_ the signal at low key b value and S_Hb_ the signal at high key b value.

Second, data were analyzed using the Sindex method [28]. The Sindex diffusion marker has been designed to identify tissue types or conditions based on their microstructure [28]. The Sindex was calculated from the direction-averaged, normalized signals, S_V_(*b*) in each voxel, as the algebraic relative distance between the vector made of these signals and those of 2 signature tissue signals S_A_ in condition A, and S_B_ in condition B, at each key b value as [28]:

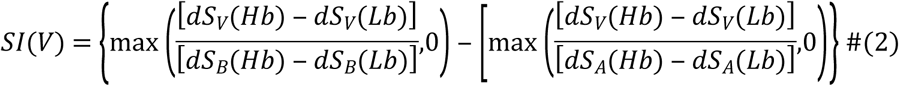

with dS_V,A,B_(*b*) = [S_V,A,B_(*b*)-S_N_(*b*)]/S_N_(*b*). S_N_ is taken as an intermediate signal between S_A_ and S_B_. SI was then linearly scaled as *Sindex* = (SI+1)*25+25 to be centered at 50. A tissue with status similar to condition A has *Sindex* = 75, while for a status similar to condition B one has *Sindex* = 25. The library of the 2 reference diffusion MRI signals was built in advance from previously acquired data, one representing a generic mouse brain tissue (B) and another derived by simulating a moderate increase in diffusion hindrance (A) using the Kurtosis diffusion model [28]. For this study, S_A_(Lb)=0.858; S_A_(Hb)=0.370; S_B_(Lb)=0.855; S_B_(Hb)=0.317. Obviously the Sindex is expected to vary widely across brain regions according to their degree of diffusion hindrance (Fig. 3), for instance white matter regions have higher Sindex values than gray matter regions. However, our focus was on the local changes in Sindex values induced by the injection of TGN-020, reflecting local changes in the degree of diffusion hindrance (a decrease in Sindex reflecting a decrease in hindrance). Sindex and sADC were calculated on a voxel-by-voxel basis to generate parametric maps. Calculation was also performed on a ROI level, averaging signals from all voxels within the ROI. ROIs were placed over the cerebral cortex, the striatum and the hippocampus (CA3 region and dentate gyrus, DG). We chose the CA3 and the DG as ROI because of abundant AQP4 expression in the DG compared with that in the CA3 [35,36].

**Fig. 3.**
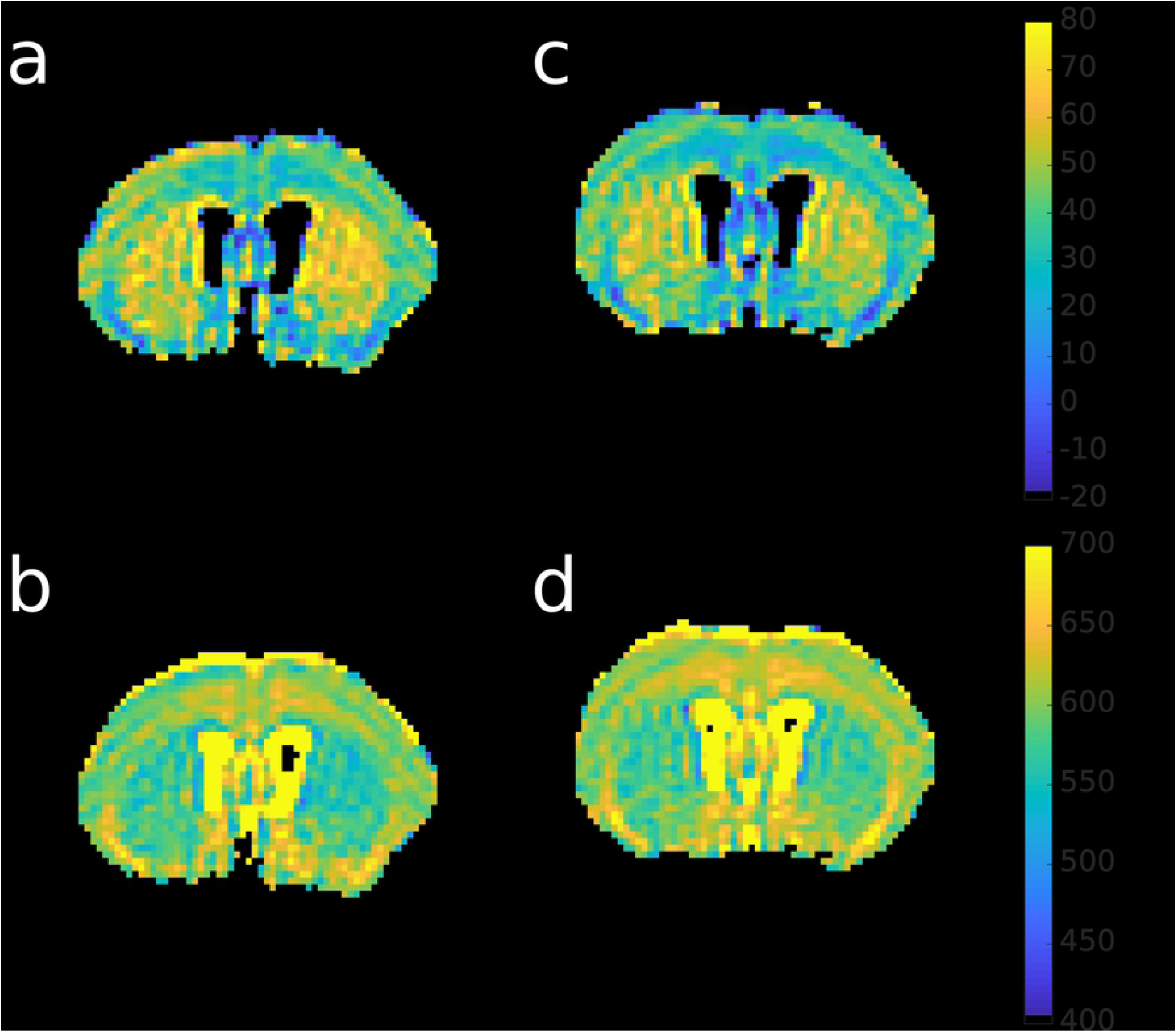
Sindex (first row) maps and sADC (second row) maps for a representative mouse from vehicle group (first column) and from TGN-020 group (right column).

Values obtained before (6 first scans) and after injection (6 last scans) were averaged in each ROI for each animal. The parameter time course was also calculated by averaging 3 successively acquired datasets (resulting in six time points with a resolution of 15 minutes) for each animal. Before averaging across individual subjects, we performed an outlier exclusion using a z-score filter (z>3) calculated over the subject’s parameter values for each time point.

#### Statistical analysis

The statistical tests were performed in python (Python Software Foundation. Python Language Reference, version 3.7. Available at http://www.python.org). We performed a paired two sample t-test between pre and post-injection data in each group and a pair-wise t-test with a posthoc correction for multiple comparison to compare the post-injection data between two groups and the pre-injection data between two groups.

For the time course data, we performed a two sample t-test between the two groups for each time point.

## Results

### Averaged Sindex and sADC following vehicle or TGN-020 injection

Figure 3 shows brain maps of Sindex and sADC for a representative mouse following vehicle and TGN-020 injection. Differences between the two conditions are readily visible with a decrease in Sindex and an increase in sADC following TGN-20 injection. Those changes are quantitatively assessed in Figure 4. Fig. 4a-d show the Sindex averaged over six scans before and after the injection in different locations, i.e., cerebral cortex, hippocampus and striatum, in vehicle and the TGN-020 group. TGN-020 injection resulted in significant decrease in Sindex in the cortex (p = 0.0061) and the hippocampus (CA3: −4.8, p = 0.00012, DG: −11.8, p = 4.8e-7) but not in the striatum (p = 0.26). No significant change of Sindex was observed following vehicle injection in those locations (p = 0.94 in cortex, p = 0.25 in CA3 and p = 0.14 in striatum), except for DG the Sindex was slightly but significantly decrease (−3.4, p = 0.03). Fig. 4e-h show the sADC averaged over six scans before and after the injection in the same locations as for the Sindex, in vehicle and TGN-020 group. TGN-020 injection resulted in significant increase in sADC in the cortex (p = 0.0064) and the hippocampus (CA3: +10.2, p = 0.00013, DG: +22.5, p = 7.14e-7) but not in the striatum (p = 0.26). No significant change of sADC was observed following vehicle injection in any of the ROIs considered (p = 0.91 in cortex, p = 0.31 in CA3 and 0.14 in striatum), except for DG the sADC was slightly but significantly increase (+6.2, p = 0.03). Interestingly, the amount of Sindex and sADC changes were different within subregions of the hippocampus, with Sindex and sADC changes much larger in DG than CA3. Those differences colocalize with known variations in AQP4 expression: AQP4 channels abundantly exist in the dentate gyrus (DG) of the hippocampus compared to other regions like CA3 region [35,36].

**Fig. 4.**
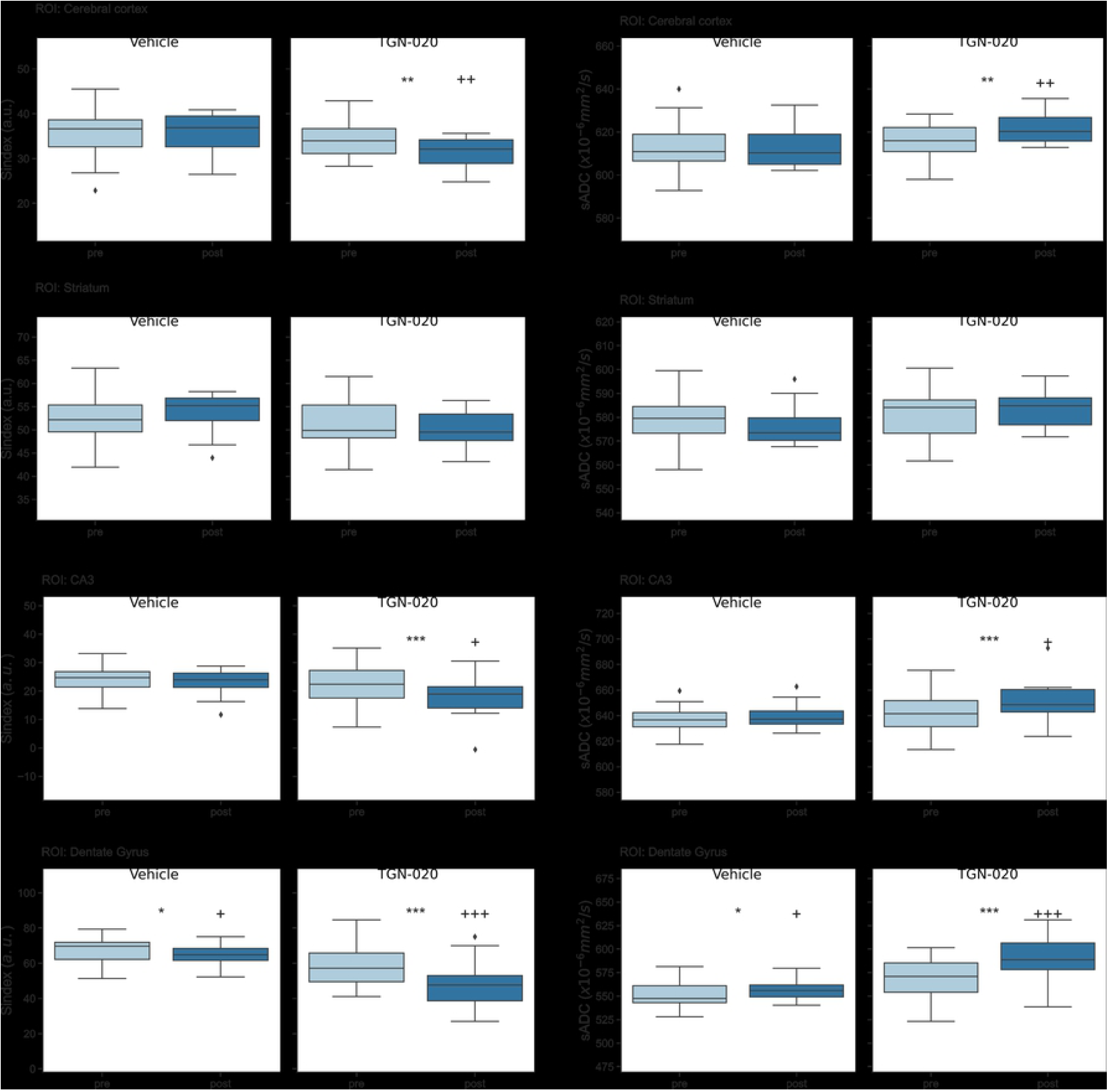
Boxplot of Sindex and sADC in the cortex (a: Sindex, e: sADC), the striatum (b: Sindex, f: sADC), CA3 (c: Sindex, g: sADC) and DG (d: Sindex, h: sADC). Left, Vehicle group; right TGN-020 group in each figure. Pre, pre-injection (light blue); post, post-injection (dark blue) of vehicle or TGN-020 group. *: p<0.05, **: p<0.01, ***: p<0.001 are the result for the paired t-test between pre and post-injection for each group. +: p<0.05, ++: p<0.01, +++: p<0.001 are the result for a t-test with posthoc correction for multiple comparison between post-injection of the two groups. Diamonds represent outliers, a point is defined as an outlier if it’s value is below Q1 – 1.5×IQR or above Q3 + 1.5×IQR, where Q1 is the first quartile, Q3 the third quartile and IQR the interquartile range.

### Sindex and sADC time courses following vehicle or TGN-020 injection

We then investigated time course of the Sindex and sADC for each group. The same ROIs were used as for averaged Sindex and sADC. Fig. 5a-d represent the Sindex time course in the different ROIs. A decrease of Sindex was observed after the injection of TGN-020 but not with vehicle injection in all ROIs. The Sindex significantly decreased (around 9% in the cortex, 20 % in DG) following TGN-020 injection in all ROIs and it continued until the end of the scanning period. Fig. 5e-h show the sADC time course with an opposite trend compared to Sindex in all ROIs.

**Fig. 5.**
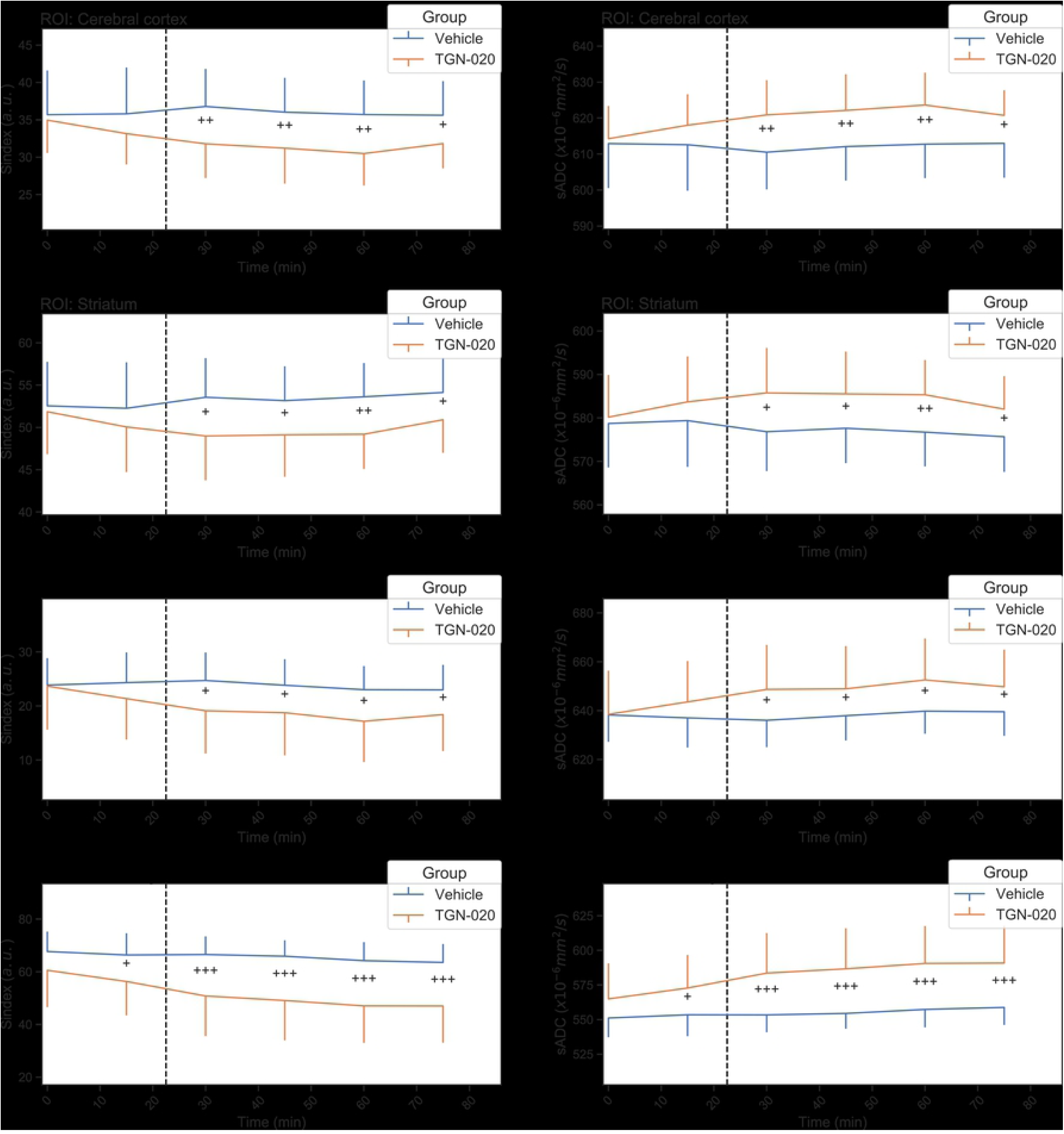
Time course of Sindex and sADC in the cortex (a: Sindex, e: sADC), the striatum (b: Sindex, f: sADC), CA3 (c: Sindex, g: sADC) and DG (d: Sindex, h: sADC). Blue line is vehicle group and orange line is TGN-020 group. Error bar shows standard deviation. The dashed line represents the injection time. +: p<0.05, ++: p<0.01, +++: p<0.001 by two sample t-test between the two groups for each time point.

## Discussion

Several studies have underlined the potential of Diffusion MRI to investigate the glymphatic system [7,22–24] and the ADC has been proposed as a biomarker of AQP gene expression, as earlier studies have demonstrated that the ADC obtained with high b values was correlated with the amount of increasing expression of AQP1 in glioblastoma cell lines [37]. In this study, we have investigated whether two diffusion MRI markers, Sindex and sADC, could detect changes in astrocyte activity induced by a pharmacological challenge. Astrocytes are considered to play a key role in brain waste clearance through membrane AQP4 channels expressed at their end-feet [12].

In this study, following AQP4 channel inhibition with a TGN-020 solution a decrease in Sindex and increase in sADC were readily observed in the cortex, more in the hippocampus, but not in the striatum, reflecting local differences in astrocyte density [38]. In the hippocampus changes in the diffusion biomarkers were larger in the dentate nucleus than in CA3 layer, presumably in line with differences in AQP4 channels expression on astrocytes [39]. Those results suggest that diffusion MRI is sensitive to astrocyte activity and, indirectly, to the status of the glymphatic system. Those diffusion MRI markers provide a higher sensitivity to small changes in tissue features by encompassing in a single marker Gaussian and non-Gaussian diffusion effects. Furthermore they are easy to calculate and are not diffusion signal model dependent, such as the kurtosis model [28]. The low key b value used in this study is high enough (250 s/mm^2^) to make perfusion-related IVIM effects negligible. Hence the observed changes in the diffusion markers reflect genuine tissue microstructure related diffusion effects and not perfusion effects (cerebral blood flow) which have already been reported observed with TGN-020 [40].

The Sindex decrease and the sADC increase jointly point out to a decrease in the amount of hindrance for water diffusion in astrocyte rich areas (cortex and hippocampus) subject acute AQP4 channel inhibition induced by TGN-20. Based on established diffusion MRI mechanisms [25] this hindrance decrease suggests an astrocyte volume reduction [41,42] and an increase of the ISF (were diffusion is tortuous) [43] resulting from the altered astrocyte function. Indeed, astrocytes rapidly regulate their volume throughout AQP4 channels. Those results obtained by acute AQP4 inhibition contrast earlier reports using chronic models, such as an ADC decrease observed during AQP4 inhibition with interfering RNA [44] or change in ADC in AQP4 knockout mice [45]. Beside the higher sensitivity to tissue microstructure of the sADC over the ADC, such discrepancy could possibly result from the modified astrocyte phenotype associated with a long-term inhibition of AQP4 expression found in those previous studies [46]. The sub-acute or chronic inhibition of AQP4 activity by AQP4-antibodies or small interfering RNA duplexes alter astrocyte morphology and decrease water permeability [47,48] which could result in an ADC decrease. Also, the effects we observed in baseline conditions should be distinguished from those obtained in conditions of neuronal activation for which AQP4 inhibition by extracellular acidification results in astrocyte swelling, capillary lumen expansion and Virchow-Robin space reduction [49]. Clearly, the detailed mechanisms in vivo underlying AQP4 channel inhibition by TGN-20 and in AQP4 knockout mice models are lacking, and diffusion MRI studies have the potential to clarify those mechanisms.

Another possible confound is that our studies were obviously performed under anesthesia using low doses of dexmedetomidine and isoflurane. Anesthetic drugs are known to impact on intracranial pressure, which could interfere with CSF-ISF exchanges. For instance, dexmedetomidine and ketamine enhance the CSF influx alongside perivascular spaces [50]. Moreover, similar to what happens during sleep, anesthesia is associated with a substantial increase in both perivascular and extracellular space volume [51]. Thus, the effects we have observed with TGN-20 induced under anesthesia might be peculiar to this this brain state and not necessarily be identical in awake animals.

## Conclusion

Work remains to better understand the mechanisms involved in brain waste clearance and the contribution of astrocytes. The reference method to investigate the so-called glymphatic system in preclinical settings relies on the intracisternal injection of gadolinium in the cisterna magna [52]. Our results show that changes in astrocyte activity, an important component of glymphatic system functionality, could be monitored non-invasively with diffusion MRI, especially through its Sindex metric. Because it is non-invasive, this approach could be used for clinical studies to confirm or infirm the existence of such a glymphatic system in humans [23].

## Acknowledgements

This research was supported by a public grant of the French National Research Agency (project “MrGLY” (reference: ANR-17-CE37-0010)).

